# Epistasis between drug resistance-conferring mutations in *Mycobacterium tuberculosis*

**DOI:** 10.1101/2025.08.07.669101

**Authors:** Selim Bouaouina, Sònia Borrell, Jérôme Azizaj, Thomas Bock, Amanda Ross, Chloé Loiseau, Sevda Kalkan, Miriam Reinhard, Valerie F. A. March, Anna Dötsch, Alexander Schmidt, Amir Banaei-Esfahani, Sebastien Gagneux

## Abstract

Studies in model organisms show that mutations conferring resistance to different antibiotics can interact epistatically, but the biological and epidemiological consequences of such interactions are unknown. Here we show that in *Mycobacterium tuberculosis (Mtb)*, positive sign epistasis between RpoB and GyrA mutations causing resistance to rifampicin and fluoroquinolone, respectively, can lead to double-resistant strains with high *in vitro* fitness. Two of these RpoB-GyrA mutation combinations account for 53% in a global collection of highly drug-resistant *Mtb* clinical isolates, compared to <0.7% for RpoB-GyrA combinations with low *in vitro* fitness. Moreover, the two high-fitness RpoB-GyrA combinations are associated with a more benign and idiosyncratic proteome perturbation compared to low-fitness combinations. Our findings highlight the relevance of epistasis for the emergence and spread of antimicrobial resistance.

## Introduction

Epistasis refers to the phenomenon where the phenotypic effect of one mutation is influenced by one or multiple additional mutations (*1*). Studies in model organisms have shown that mutations conferring resistance to antibiotics can interact epistatically, sometimes resulting in bacterial genotypes resistant to two drugs showing a higher fitness than the corresponding genotypes resistant to only one drug (*2–5*). However, the biological and epidemiological consequences of such epistatic interactions have not been studied in any obligate human pathogen. *Mycobacterium tuberculosis (Mtb)*, the etiological agent of human tuberculosis (TB), remains the number one cause of human death from a single infectious agent (*6*). The standard treatment for TB comprises a combination of multiple antibiotics administered over six months (*7*). Multidrug-resistant TB (MDR-TB) is defined by resistance to isoniazid and rifampicin, the two most important first-line drugs used to treat drug-susceptible TB (*6*). MDR-TB is treated with second-line agents that include fluoroquinolones. *Mtb* strains resistant to both first- and second-line drugs are a growing public health concern (*6, 8*). As *Mtb* shows no evidence of ongoing horizontal gene exchange (*9*), all drug resistance-conferring mutations are encoded chromosomally and acquired sequentially (*10*). Compensatory mutations in the bacterial RNA polymerase have been shown to enhance the fitness of highly drug-resistant *Mtb* clinical strains *in vitro* (*11*), during patient-to-patient transmission (*12*), as well as within patients, thereby enhancing the odds of acquiring additional drug resistance mutations (*13–15*). However, if and how the various drug resistance mutations interact epistatically to influence the fitness of highly drug-resistant *Mtb* clinical strains has not been addressed.

Here, we tested for epistasis between RpoB and GyrA mutations known to confer high-level resistance to rifampicin and fluoroquinolones, respectively. We compared the *in vitro* fitness of a set of defined *Mtb* mutants that harbored a resistance-conferring mutation in one or both of these genes. We then compared our *in vitro* data to the clinical frequency of the corresponding double-mutants using whole genome data from a global collection of 109,332 *Mtb* clinical strains. Finally, we explored the phenotypic consequences of these epistatic interactions using comparative proteomics.

## Materials and Methods

### Selection of strain set and growth conditions

Of a previously described and characterized set of pan-susceptible MTBC clinical isolates (Borrell, 2019), we selected three strains each of genetically distinct sub-lineages of MTBC Lineage (L)1, L2 and L4 (Table S1). In our study, we refer to genetic traits shared between strains of the same lineage and genetic traits specific to each strain separately as *lineage-background* and *strain-background*, respectively. The strains were grown in 50mL conical tubes (Falcon Corning, NY, USA) shaking (37°C, 140rpm, 2.5cm swing) in Middlebrook 7H9 broth, supplemented with 10% ADC (7H9 ADC, 5% bovine albumin-fraction V + 2% dextrose + 0.003% catalase, Sigma-Aldrich), 0.5% glycerol (PanReac AppliedChem) and 0.05% Tween-80 (Sigma-Aldrich). Solid cultures were grown on Middlebrook 7H11 agar, supplemented with ADC and oleic acid. All handling of infectious material was conducted in a biosafety level 3 laboratory.

### *In vitro* spontaneous mutant selection

Exponentially (OD_600_=0.6±0.1) growing *Mtb* liquid cultures were spun down (800xg, 5min, 4°C), the supernatant removed, the pellet resuspended in 300uL 7H9-ADC and plated on 7H11-OADC with 5ug/mL Rifampicin (Sigma-Aldrich) and/or 2ug/mL Ofloxacin (Sigma-Aldrich). Single colonies were picked and regrown in 7H9-ADC. When in late exponential phase, three 1mL aliquots were taken from each culture, two of them frozen at −80°C and one of them heat-inactivated (95°C, 1h) for PCR of the genes of interest, *rpoB* (forward primer: TCGGCGAGCTGATCCAAAACCA; reverse primer: ACGTCCATGTAGTCCACCTCAG) and *gyrA* (forward primer: CAGCTACATCGACTATGCG; reverse primer: GGCTTCGGTGTACCTCATC). The amplicon was Sanger-sequenced at Microsynth (Balgach SG, CH), and the sequences analyzed with Ugene v50.0 (*38*). Frozen aliquots of the samples carrying the mutation of interest (RpoB S450L, RpoB H445R, GyrA D94G, GyrA A90V, GyrA G88C alone and in combination) were regrown in 7H9-ADC and molecular DNA extracted following the previously described CTAB method (39). The DNA was then paired-end sequenced on an Illumina HiSeq 2500, Ilumina NovaSeq 6000 and Illumina MiSeq sequencing platform at the genomics core facility of the ETH Zürich and University of Basel in Switzerland. The average per-run read length was 118bp (from 38-153), with a resulting average sequencing depth of 132 (11–383). We deposited the data in the European Nucleotide Archive using the following project accessions PRJEB94519 (see Data S1 for individual accession numbers). All Illumina reads were analyzed with our in-house analysis pipeline as described previously (*15, 19*) and the genomes screened for potential off-target mutations.

### Off-target variant identification and SNP distance calculation

All Illumina reads were processed and analyzed as described in (*15, 19*). We identified off-target variants, which were variants with an allele frequency ≥90% in a mutant strain but not present in the respective wild type ancestor, by comparison of the respective VCF files. For SNP distance calculation between all strains, we used *snp-dist* (GitHub - tseemann/snp-dists: Pairwise SNP distance matrix from a FASTA sequence alignment) with default settings (Data S3).

### *In vitro* growth curves

Exponentially growing *Mtb* cultures were calibrated to OD_600_=0.1 and triplicates inoculated with 250uL, each 15mL drug-free 7H9-ADC, containing 2.5mL equivalent of 2mm borosilicate beads. Starting 48h post inoculation, growth of each sample was measured twice a day by 10 seconds of vortexing to disrupt aggregation, and measuring optical density at 600nm (OD_600_), using 60uL of culture in UVette® (Eppendorf®, Hamburg, Germany) in an Ultrospec™10 (Harvard Biochrom, Holliston, MA, USA).

### Estimating the relative fitness

To account for measurement errors by the spectrophotometer or due to cellular aggregation at increased culture density, the measured data was filtered for 0.03 ≤ OD_600_ ≤ 0.70 prior to log2- transformation. To estimate fitness, we first identified the subset of data points for each replicate, which corresponded to the growth rate uninhibited by culture density. We did so by fitting linear models for each possible combination of at least six consecutive observations and selecting the linear model with the steepest slope and an 𝑅^2^ ≥ 0.95. We estimated the relative fitness of each single or double mutant strain, as the ratio of the respective Malthusian parameters, 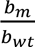, where 𝑏*_m_* is the slope of the mutant strain and 𝑏*_wt_* the slope of the cognate wild type strain. The slopes were estimated using a linear regression model for each mutant or wild type strain with coefficients for the intercept and slope and random effects for slope and intercept to account for the clustering of observations within replicates. We fitted the model into a Bayesian framework using the Hamiltonian Monte Carlo algorithm in Stan (*40*) via the R (*41*) interface package RStan (*42*). We used 2,000 iterations and four chains. To compare the relative fitness between the single- and double-mutants, we calculated the posterior distribution of the ratio. The full reproducible code is available on Gitlab (TBRU / mtuberculosis_epistasis· GitLab).

### Calculating the expected relative fitness of double-mutants and the corresponding epistasis

*rpoB* and *gyrA* encode subunits of very central and supposedly co-dependent proteins, active in the transcription and replication. Therefore, the expected relative fitness values for the double-mutants, were calculated by multiplying the relative fitness of the respective single RpoB and GyrA single mutants (*1, 43*). Intergenic epistasis between RpoB and GyrA was calculated as the deviation of the *measured* double mutant fitness from the *expected* double mutant fitness. The Bayesian framework allowed us to incorporate the uncertainty in the estimated relative fitness values into the estimates of the group relative fitness and intergenic epistasis.

### Associating genetic traits with relative fitness

We assessed the proportion of variance in the estimated relative fitness values that was explained by each variable by comparing models with and without each variable. The variance of the fitted values predicted by a model provides an estimate of the variance explained by the model (Table 1).

**Table 1.** Variation in relative fitness explained by individual and combined effects of mutation, strain- and lineage-genetic background. Percentage of the total variance in relative fitness explained by models with different combinations of explanatory variables (M = drug resistance mutation, S = strain-genetic background, L = lineage-genetic background). + = main effects only; * = main effects and interaction terms

|  | Independent variables and their relationships | Median | Lower 95% CI | Upper 95% CI |
| --- | --- | --- | --- | --- |
| Full model | relative fitness ~ M * (L + S) | 98.6% | 95.6% | 99.6% |
| Lacking one interaction | L + M * S | 98.6% | 95.6% | 99.6% |
|  | S + M * L | 84.8% | 79.6% | 89% |
| No interaction | S + M + L | 66.7% | 60% | 72.5% |
| Lacking one or two variables | M * S | 98.6% | 95.6% | 99.6% |
|  | M * L | 72.3% | 66.4% | 77.5% |
|  | S * L | 25.0% | 18.4% | 31.8% |
|  | L | 1.2% | 0.1% | 3.8% |
|  | S | 25.0% | 18.4% | 31.8% |
|  | M | 45.8% | 39.9% | 51.7% |

### Bioinformatic analysis of *Mtb* genome data from the clinic

From 109,332 publicly available good quality genomes (mostly coverage ≥ 20X, but at least ≥ 10X after analysis according to the same method described above), we removed the mixed infections (ratio variable SNPs to fixed SNPs ≤ 0.2) and selected for MTBC L1-L10, ending up with 90,144 genomes. Of these 90,144 genomes, we filtered for pre-XDR (resistant to rifampicin and a fluoroquinolone) and XDR (additional resistance to Bedaquiline or Linezolid) genomes, which carry one or multiple known drug resistance mutation in *rpoB* and *gyrA* at an allele frequency of >= 10% and ended up with 9,096 genomes. If the genome has been sequenced in Western-Europe, USA or Canada, we used the indicated patient’s birth place, if the genome was sequenced elsewhere, we used the respective country to determine where the sample originates from. We ended up with genomes stemming from 50 countries covering five human-adapted lineages of the MTBC (L1-L4, L6).

### Geographical maps

A geographical world map was loaded from the R-package *rnaturalearth* (*44*) and per-country centroids from the GitHub repository curated by Gavin Rehkemper (https://github.com/gavinr/world-countries-centroids), freely available under an MIT license.

### Full proteome extraction

*Mtb* strains were grown in 10mL drug-free 7H9-ADC to exponential phase, calibrated to an OD_600_=0.1 and 1mL inoculated in quadruplicates of 100mL 7H9-ADC each, supplemented with 10mL equivalent 2mm borosilicate beads. Exponential phase-cultures were washed twice with cold PBS (3500xg, 6min, 4°C), and the dry pellet snap frozen in liquid nitrogen and stored at minus 80°C. For proteome extraction, the frozen pellets were resuspended in 500uL diluted lysis buffer (0.1% SDC, 10mM TCEP, 100mM Tris) and the cellular proteins denatured for 10min at 95°C. The suspension was then transferred to bead-beating tubes (MP Biomedicals, CA, USA) containing 500uL equivalent of 0.1mm silica spheres. The samples were then mechanically disrupted for 4x30s at 9,000rpm on a homogenizer (Precellys Evolution, Bertin Technologies SAS, Montigny-le-Bretonneux, F), with 1min cooling on ice between the bead beating. The tubes were then spun down (5min, 13,000xg, 4°C) and the supernatant with the extracted proteins was transferred to another tube and 500uL of diluted lysis buffer added to the bead-beating tubes, followed by one cycle of homogenizing. The SDC concentration in the supernatant was increased to 1% (by adding concentrated lysis buffer, 10% SDC, 10mM TCEP, 100mM Tris). The tubes were then incubated on ice for 15min before being filter-sterilized twice through 0.22um porous filters mounted on syringes (BD AG, Allschwil, CH). Samples were extracted in batches of 15 in a randomized order.

### Sample preparation for MS-based proteome analysis

Samples were heated to 95° C for 10min and sonicated on a Pixul® Multi-Sample Sonicator for 20 min. The protein concentration was adjusted to 50 ug per sample and carbamidomethylation was performed by addition of Chloroacetamide (Sigma, C0267-100G, final concentration 15 mM). Samples were digested by incubation with 1 ug Trypsin (Promega, V5113) at 37°C overnight. Tryptic activity was stopped by addition of 5% TFA and peptides were washed and purified using iST cartridges (Preomics.com) according to the protocol of the manufacturer. Peptide samples were dried by vacuum concentration and stored at −20° C until further use. All samples were dissolved in LC-buffer containing 0.1% FA/in H2O prior to the injection into the mass spectrometer.

### Mass Spectrometry Analysis

For each sample, 0.3 µg total peptides were subjected to LC-MS analysis using Sample Block Randomization on a Orbitrap Exploris 480 mass spectrometer equipped with a nanoelectrospray ion source (both Thermo Fisher Scientific). Peptide separation was carried out using an Evosep One chromatographic system (Evosep.com) equipped with a Performance column (150 μm x 15 cm, 1.5 um, EV1137) and a custom-made column heater (50° C) at 500 nl/min flow rate (30 SPD method). The Orbitrap Exloris 480 mass spectrometer was operated in data independent acquisition mode (DIA) with an approximate cycle time of 3 seconds. Each cycle, a MS1 scan was acquired in the Orbitrap in centroid mode at a resolution of 120,000 FWHM (at 200 m/z), a scan range from 350 to 1500 m/z, normalized AGC target set to standard and maximum ion injection time mode set to “auto”. A total of 42 DIA scans were acquired in the Orbitrap per cycle (centroid mode) at a resolution of 15,000 FWHM (at 200 m/z), a precursor mass range of 400 to 904, quadrupole isolation window of 12 m/z, a defined first mass of 120 m/z, normalized AGC target set to 3000% and a maximum injection time set to “auto”. Peptides were fragmented by HCD (Higher-energy collisional dissociation) with normalized collision energy set to 28 % and one microscan was acquired for each spectrum. Generated raw files were searched against the *Mycobacterium tuberculosis* database (ATCC25618, H37Rv, UP000001584, downloaded from Uniprot 20230923, 3,996 proteins) and 392 commonly observed contaminants using using SpectroNaut (Spectronaut 19.0.240606.62635, Biognosys) with default settings.

### Full proteome analysis

The data was then submitted to our automated R-based in house proteomic analyses pipeline “tbeeprotpip” (https://git.scicore.unibas.ch/TBRU/tbeeprotpip) where outlier runs were removed, protein intensities were normalized by median, followed by removal of non-*Mtb* proteins using functions from MSstats (*45*) and protti (*46*). Differentially abundant proteins (DAP) between samples were calculated using MSstats. For the differential abundance analysis, we conducted multiple sensitivity analyses and found these results to be robust to different levels of statistical significance and FC-thresholds (Table S3). In a next step, enrichment analysis, was conducted as described before (*47, 35*) using clusterProfiler (*48*). The full reproducible code is available on Gitlab (TBRU / mtuberculosis_epistasis · GitLab) and the initial table with the proteome data is on Zenodo (doi: 10.5281/zenodo.16744470).

### Associating genetic and proteomic distances

SNP distances between samples (see above) were associated with proteomic distances (Euclidean distance) between samples using a Mantel test from the *ade4* R-package (*49*).

## Results

### Isolating spontaneous single- and double-resistant mutants

We isolated spontaneous single- and double-mutants with amino acid substitutions in RpoB and/or GyrA known to confer a high level of resistance to rifampicin and fluoroquinolones, respectively. These spontaneous mutants were generated starting from nine drug-susceptible *Mycobacterium tuberculosis* complex (MTBC) clinical strains, henceforth referred to as wild type strains. These nine wild type strains included three phylogenetically distinct strains, each from one of the three MTBC lineages L1, L2 and L4 (*16*). While L1 has been associated with lower virulence in infection models (*17, 18*), reduced transmission between patients (*19–22*), and low levels of drug resistance, the opposite is true for L4 (*23*) and particularly L2 (*12, 24, 25*). Because multiple mutations in RpoB and GyrA can cause drug resistance (*26*), we focused on mutations previously shown to be individually associated with either a low- or a high fitness cost *in vitro* (*27, 28*). Specifically, RpoB S450L, GyrA D94G and GyrA A90V were considered low fitness cost mutations, and RpoB H445R and GyrA G88C high fitness cost mutations. Starting from the nine wild type ancestor strains, we isolated 40 out of 45 (89%) combinatorially possible single RpoB and GyrA mutants carrying any of these five mutations, and 34 out of 54 (63%) possible double-mutants (Table S1). The fact that we failed to isolate all possible mutants in our matrix, despite multiple attempts, indicates synthetic lethality in the mutants we were unable to generate (i.e. they were not viable). Whole genome analysis of the 74 different mutants we managed to isolate, revealed a total of 16 unique off-target mutations (Table S2). Eleven of the 16 unique off-target mutations occurred in only one strain each. Of the 74 mutants analyzed, 53 (72%) harbored no off-target mutation. Seven single- and 13 double-mutants each carried one off-target mutation. One mutant carried two off-target mutations. Two known and two potential new compensatory mutations in the RNA polymerase were detected in seven double-mutants, but not in any of the single-mutants.

### The fitness effects of RpoB and GyrA mutations are strain-specific

We calculated the relative fitness based on the *in vitro* growth of each single- and double-mutant within each strain-background, across strain-backgrounds, and for each wild type strain compared to the wild type strain L4_C00 (see Methods; Fig. 1, S1 & S2, Data S1). Across all strain-backgrounds, GyrA D94G mutants showed a negligible fitness cost, with the other GyrA single-mutants, the RpoB S450L single-mutants and the double-mutants carrying low fitness cost RpoB and GyrA mutations exhibiting a fitness cost of <10%. By contrast, the RpoB H445R single-mutants and all double-mutants carrying RpoB H445R and/or GyrA G88C exhibited a higher fitness cost (>20%). When comparing the wild type strains between each other, we found that most wild type strains were similar but L1_A00 had a lower relative fitness. When comparing the relative fitness of all single-mutants to all double-mutants, we found that, on average, the single-mutants had a higher fitness than the double-mutants (single-to-double-mutant fitness ratio of 1.09 [95% CI: 1.06, 1.11]). This indicates that acquiring resistance to an additional drug results in a further reduction in fitness.

**Fig. 1.**
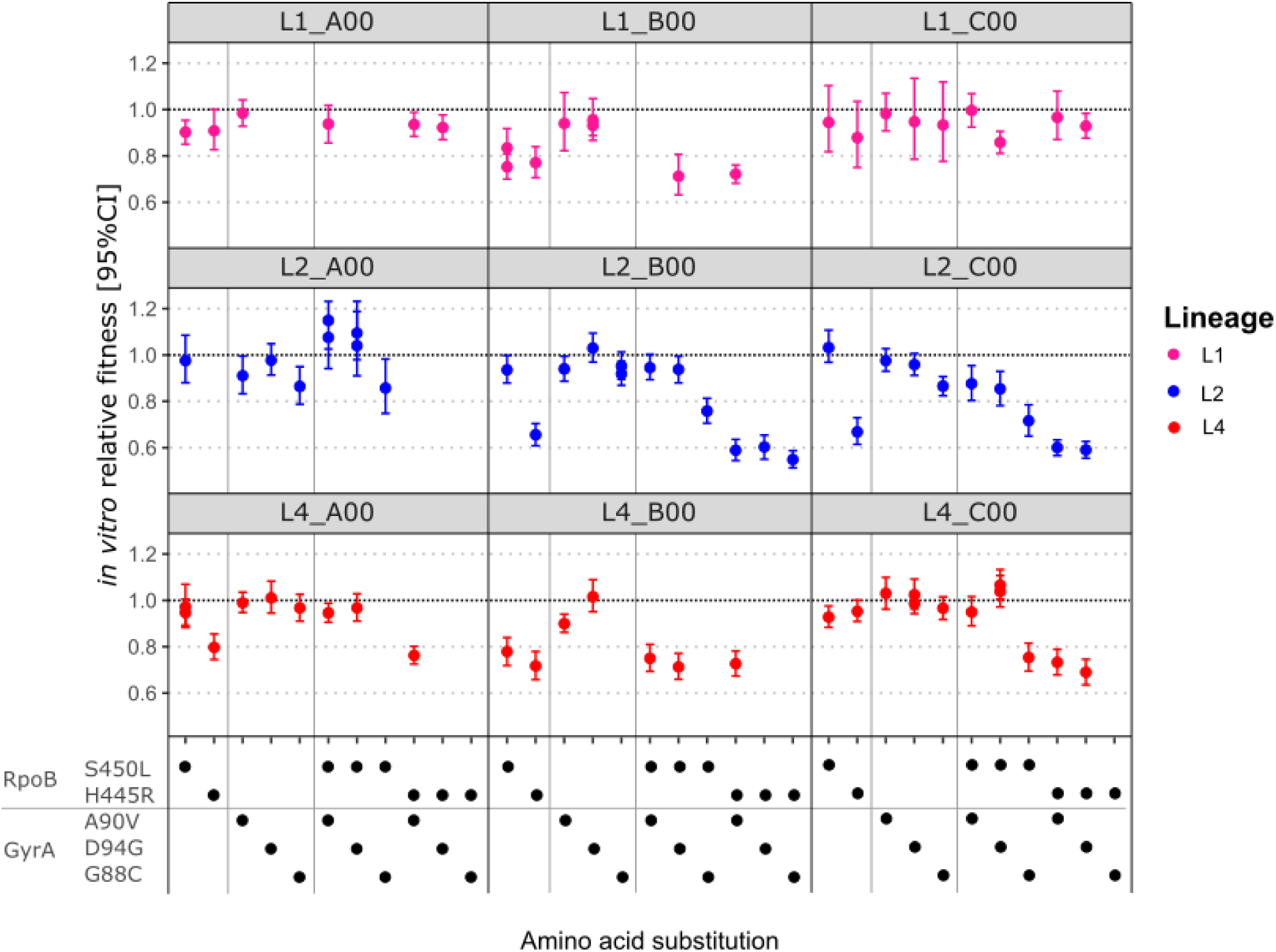
In vitro relative fitness of single- and double-mutants of the nine wild type strains. Horizontal dashed black line = fitness of the respective wild type strain.

Next, we assessed the relative contribution of drug resistance mutations alone and in combination, and the strain- and lineage-backgrounds in explaining the variation in relative fitness values using a multivariable model (Methods, Table 1). We found that the drug resistance mutation had the largest impact overall, while the effect of the lineage-background was negligible. Including interaction terms in our model supported epistatic interactions between drug resistance mutations and strain-background (Table 1). In summary, our findings showed that the relative fitness of our single- and double- drug-resistant mutants was heterogeneous and strain-specific and mostly influenced by the specific drug resistance mutations and the strain-background.

### Epistasis between RpoB and GyrA drug resistance mutations *in vitro*

To formally test for epistatic interactions between the different drug resistance mutations in RpoB and GyrA, we calculated the expected relative fitness of the respective double-mutant, using a multiplicative model (see Methods). For every RpoB and GyrA mutant-pair in each strain background, we calculated the pairwise epistasis (Ɛ), as the deviation of the observed from the expected relative fitness of the double-mutant. In 22/33 cases (67%), we could not detect any epistatic interaction between the RpoB and GyrA mutations (Fig. 2A & Fig. S3). By contrast, 8/33 (24%) cases showed negative epistasis, where the double-mutant had a lower fitness than expected based on the fitness of the respective single-mutants (Fig. 2B). In 3/33 (9%) cases, we saw positive epistasis, where the double-mutant had a higher fitness than expected based on the fitness of the respective single-mutants; this involved one strain belonging to L2 and two L4 strains (Fig. 2C). Furthermore, in two of these cases, we saw positive sign epistasis (*29*), where the double-mutant had a higher fitness than at least one of the corresponding costly single-mutants. This latter finding suggests that in these cases, acquiring resistance to a second drug compensated for the fitness loss associated with drug resistance to the other drug. In summary, we found that one third of the RpoB-GyrA-double-mutants showed negative or positive epistasis. Notably, we observed two instances of positive sign epistasis, one of which occurred in an L4 strain that also carried a compensatory mutation, and the other in an L2 strain without any off-target mutation.

**Fig. 2.**
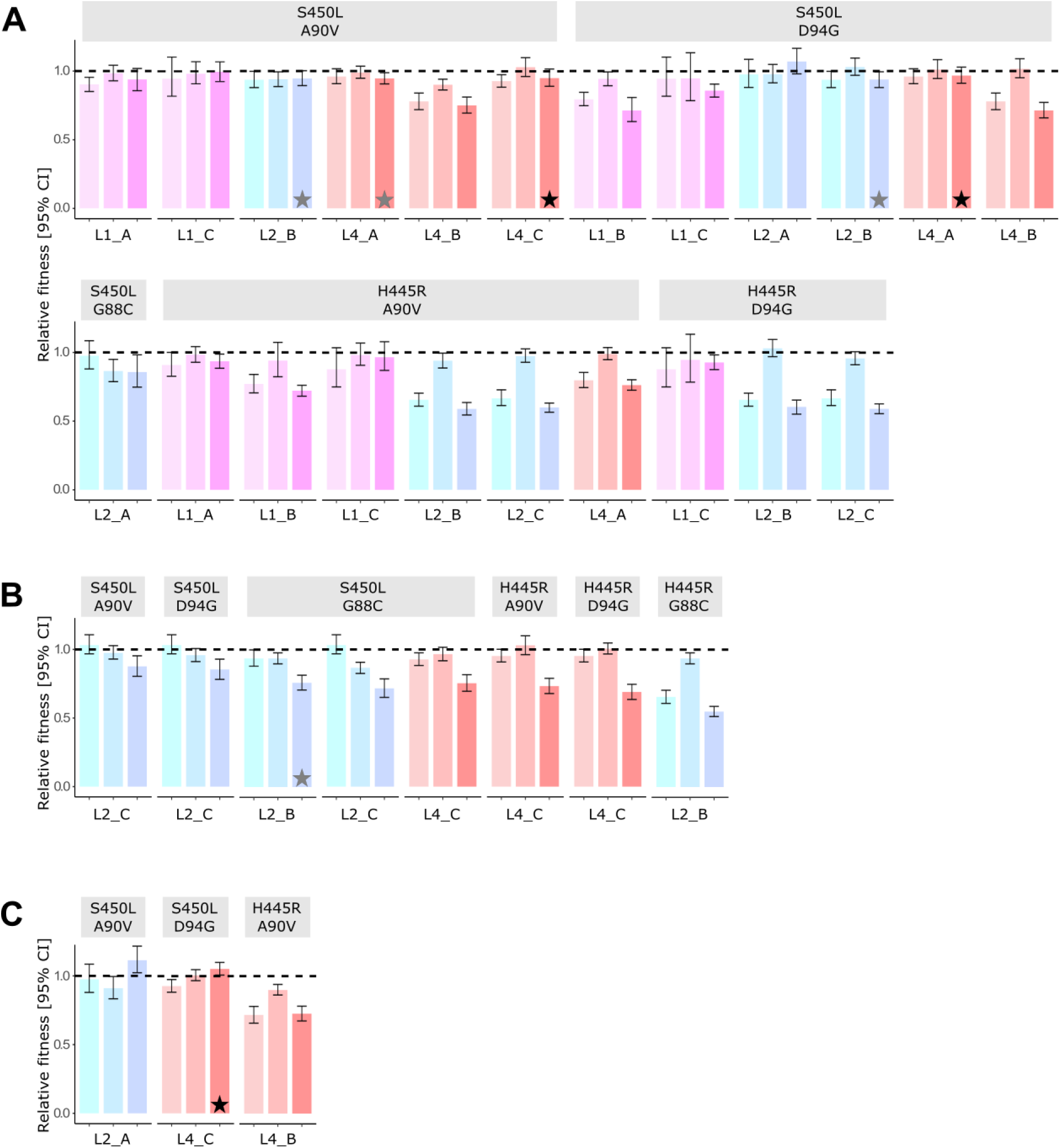
Epistatic interactions between RpoB and GyrA drug resistance mutations. (**A**) Cases where no epistasis could be detected, (**B**) cases with negative epistasis, and (**C**) cases with positive epistasis. Each colored bar shows the relative fitness of a single or double-mutant compared to its cognate drug-susceptible wild type. For each strain (x-axis labels), the RpoB and GyrA single-mutants (left and center bar, respectively) and the respective double-mutant (right bar) are shown, with 95% credible intervals. Respective RpoB (top) and GyrA (bottom) mutations are indicated in the gray bars. **★** known and **★** potential compensatory mutations.

### Positive sign epistasis between RpoB and GyrA mutations is linked to a high clinical frequency

To put the epistatic interactions between RpoB and GyrA mutations we observed *in vitro* into a clinical context, we screened 109,332 publicly available MTBC genomes and identified 8,067 genomes that carried at least one RpoB and one GyrA mutation known to confer drug resistance (*26*). According to the latest WHO definition, these genomes represent cases of pre-extensively drug-resistant (XDR) or XDR-TB (*6*). These 8,067 pre-XDR/XDR genomes originated from 50 countries across five continents and encompassed five of the ten main MTBC lineages (i.e., L1-L4, L6), which together are responsible for >95% of all TB cases in the world (*30, 31*) (Fig. S4, Data S2). These 8,067 genomes harbored 297 different combinations of fixed RpoB and GyrA drug resistance mutations, with 97% of these individually accounting for less than 1% of all the pre-XDR/XDR cases in the dataset. By contrast, 34% and 19% of these genomes carried the combinations S450L-D94G and S450L-A90V, respectively, together accounting for 53% of all pre-XDR/XDR cases in our clinical dataset (Fig. 3A). This clinical predominance is consistent with the high *in vitro* fitness of the corresponding double-mutants that also showed positive sign epistasis (Fig. 3B). Conversely, the combinations with a low *in vitro* fitness individually accounted for ≤0.7% of clinical cases.

**Fig. 3.**
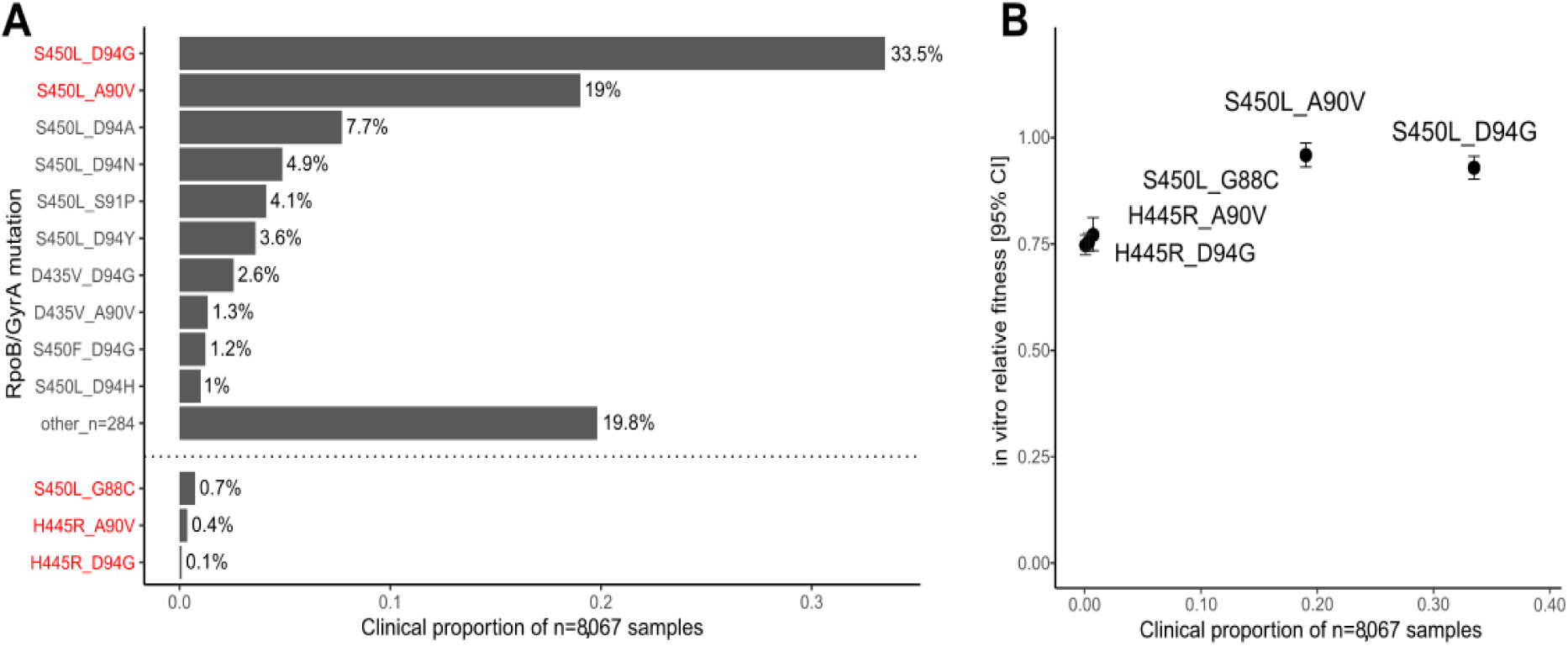
Clinical frequency of RpoB-GyrA drug resistance mutations in combination. (**A)** The clinical frequency of each combination of fixed RpoB and GyrA drug resistance mutation, which occurred in the data set of clinical isolates (n=8,067) is depicted. (**B)** Association of the *in vitro* relative fitness of the double-mutants with their respective clinical frequency. Red=the mutation combinations from our *in vitro* data, occurring in the clinic.

### The role of compensatory evolution

Because drug-susceptible TB is treated with a first-line regimen that includes rifampicin and MDR-TB is treated with second-line drugs that include fluoroquinolones, resistance to rifampicin usually arises before fluoroquinolone resistance (*32–34*). Consequently, compensatory mutations in the RNA polymerase of *Mtb* linked to rifampicin resistance will likely precede, accompany, or facilitate the emergence of resistance to other drugs including fluoroquinolones. This notion is supported by the observation that compensatory mutations have been linked to drug resistance amplification in the clinic (*13–15*). Because seven of our *in vitro*-selected double-mutants also acquired a compensatory mutation during the selection process, we determined the frequency of compensatory mutations across the different combinations of RpoB-GyrA drug-resistant mutants in our clinical genome dataset. We found that 3,098/4,236 (73%) of the high-fitness double-mutants S450L-D94G and S450L-A90V carried a compensatory mutation compared to 1,617/3,831 (42%) of the strains harboring other combinations of RpoB and GyrA mutations (OR=3.73, CI_95%_: 3.39-4.10, p<0.001). The odds for compensation did not differ between the group carrying S450L-D94G and S450L-A90V (OR=0.880, CI_95%_: 0.760-1.02, p=0.08). In summary, we found that clinically by far the most frequent pre-XDR/XDR *Mtb* genotypes carry one of the two combinations of RpoB and GyrA mutations associated with positive sign epistasis *in vitro*, often together with a compensatory mutation. Taken together, these findings suggest a role for multidimensional epistasis in the emergence and spread of highly drug-resistant *Mtb*.

### Clinically frequent and rare RpoB-GyrA variants differ in their proteome perturbation

We previously reported that five different *Mtb* clinical strains carrying the same RpoB S450L resistance mutation alone exhibited strain-specific gene expression patterns (*35*). Based on these findings, we explored the functional consequences of epistasis between RpoB and GyrA drug resistance mutations by comparing the intracellular proteome of our wild type and respective single- and double-mutant strains *in vitro* using mass spectrometry based proteomics on Data-independent acquisition (DIA) mode (Table S1, Data S1). Bacterial cultures were grown to mid-log phase in four replicates in standard Middlebrook 7H9 broth, total proteins extracted and analyzed by LC-MS/MS (see Methods). We quantified on average 3,182 *Mtb*- proteins per sample, which corresponds to 80% of the ∼4000 open reading frames annotated in the *Mtb* genome, with replicates that were of good quality (all CVs <12%, Data S1). We found that the proteomic profiles of our mutants reflected their phylogenetic relationships to large extent (see Methods, Mantel test: 999 permutations, R=0.846, p-simulated=0.001). However, mutants that carried RpoB H445R alone or in combination with any GyrA mutation clustered separately from all other wild type and mutant strains in each of the strain-backgrounds (Fig. S5), suggesting a more pronounced and shared proteome perturbation in mutants carrying RpoB H445R. We next compared the number of significantly differentially abundant proteins in all double-mutants carrying the two RpoB-GyrA combinations associated with a high clinical frequency compared to the corresponding double-mutants carrying the low-fitness RpoB H445R mutation. Across all genetic backgrounds, double-mutants with RpoB H445R consistently exhibited a greater number of shared higher and less abundant proteins than those carrying RpoB S450L (Fig. 4, Table S3). Significantly differentially abundant proteins did not share any common function across strains in either of the two mutant groups; the double-mutants carrying RpoB S450L or the double-mutants carrying RpoB H445R (Table S4). In summary, RpoB H445R induces a broader and more consistent proteomic response across strain-backgrounds, which is in contrast to the strain-specific effects observed for RpoB S450L. How this phenomenon of shared *versus* idiosyncratic proteomic perturbation might be linked to the differences in fitness between these drug-resistant mutants remains to be determine

**Fig. 4.**
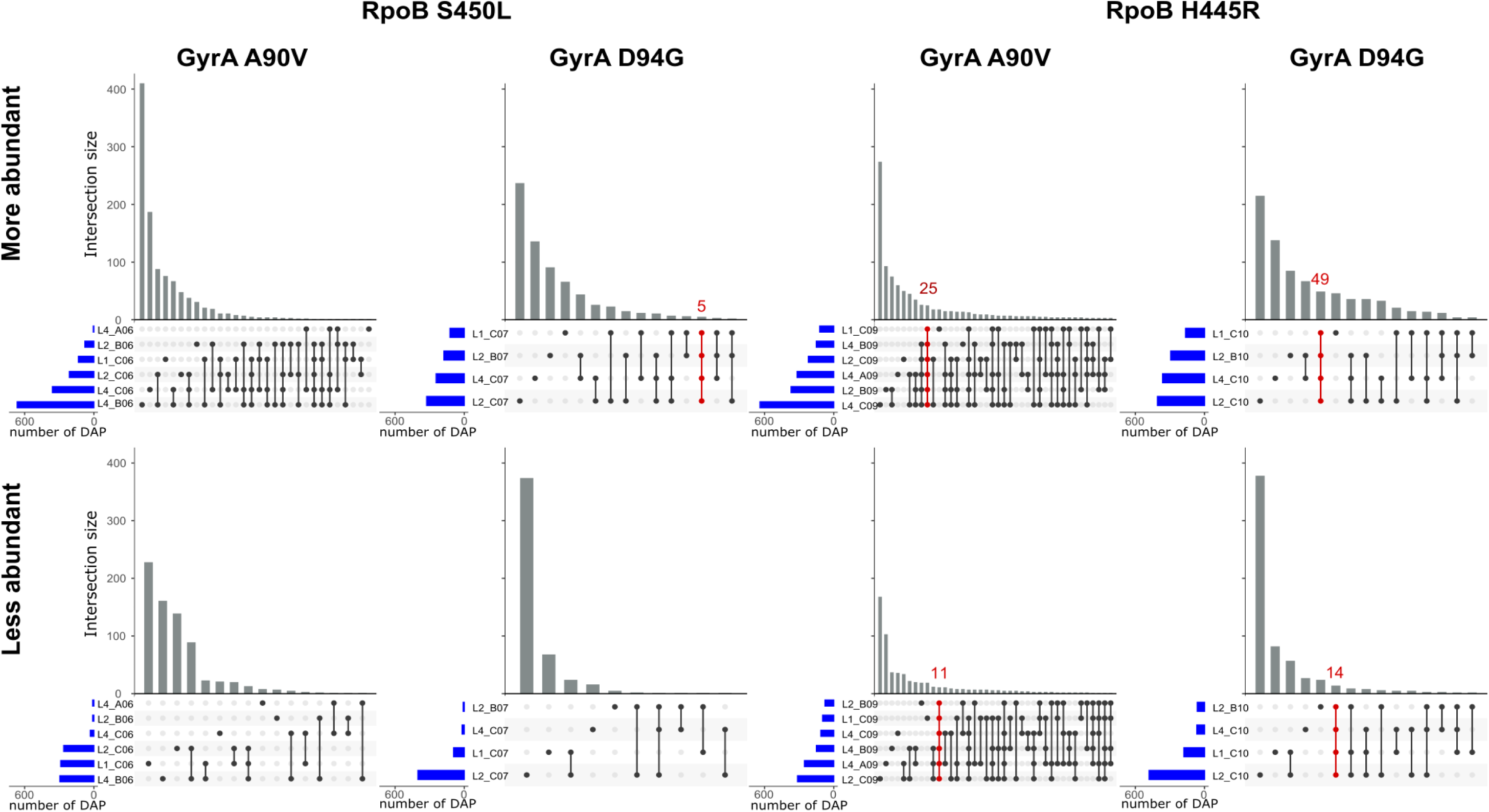
Number of significantly differentially abundant proteins (DAP) in low-fitness cost (S450L-A90V/D94G) and the cognate high-fitness cost (H445R-A90V/D94G) double-mutants. Blue bars = per-sample number of DAP; gray bars = number of DAP specific to a single sample or shared across different samples (indicated in dotplot below x-axis); red bars = number of DAP shared between all samples in one group.

## Discussion

Our study shows that drug resistance-conferring mutations in RpoB and GyrA of *Mtb* can interact epistatically. These findings are consistent with observations in other bacteria (*2–5*). Moreover, we found that the specific combinations of RpoB and GyrA mutations that exhibited positive sign epistasis *in vitro* were strongly overrepresented among drug-resistant clinical strains, indicating that the fitness of *Mtb* strains harboring such high-fitness combinations is also enhanced in the clinic. While in L1 mutants we observed no instances of epistasis between RpoB and GyrA mutations, positive epistasis occurred in three double-mutants of L2 and L4, offering a possible explanation for why L2 and L4 genotypes, but no L1 genotypes, have been associated with MDR-TB outbreaks (*31*). Finally, we found that differences in proteome perturbation might be at the basis of the fitness differences between the various drug-resistant mutants analyzed here (*35*).

Our study is limited in that the fitness assays were carried out *in vitro* and we only considered the exponential phase of bacterial growth. Yet, the parallels between the *in vitro* fitness and the corresponding clinical frequency of these mutants, indicates that our *in vitro* system is a valid proxy for *in clinico* fitness. Another limitation was the presence of some off-target mutations, revealed by whole-genome sequencing, including some instances of compensatory evolution. However, compensation could be accounted for during our down-stream analyses. Although we could not exclude an effect of compensation in all instances, compensatory mutations only emerged in 7 of 34 (20%) of our double-mutants that showed neutral, negative, and positive epistasis, and thus cannot on their own explain the full range of fitness effects observed in our study. On the other hand, our epidemiological findings showed that the majority of *Mtb* genomes carrying any of the two high-fitness RpoB/GyrA mutation combinations also harbored compensatory mutations. This supports the view that epistasis operating in this context is likely multidimensional, not only involving the different drug resistance mutations, but also compensatory mutations as well as the specific strain backgrounds.

In conclusion, our findings highlight the role of epistasis in the evolution of multidrug resistance in *Mtb*. Given the high degree of conservation of RpoB and GyrA across bacteria (*36, 37*), our findings are relevant for other multidrug-resistant bacterial pathogens.

## Acknowledgement

Calculations were performed at sciCORE (http://scicore.unibas.ch/), the scientific computing core facility at the University of Basel. The sequencing was carried out at the genomics core facility of the University of Basel and the Department of Biosystems Science and Engineering at ETHZ in Basel, Switzerland.

## Author Contributions

Conceptualization: SBou, SBor, SG. Methodology: SBou, SBor, TB, AS, SG. Investigation: SBou, JA, TB, SK, SBor, VFAM, MR, AD, AB. Formal analysis: SBou, TB, AR, CL. Data curation: SBou, JA, TB, CL, SBor, AB. Visualization: SBou, AB. Software: SBou, CL. Funding acquisition: SG. Project administration: SBou, SBor, SG. Supervision: SBou, SBor, SG, TB, AS. Writing: SBou, TB, AR, SBor, SG. Writing – review & editing: SBou, SG, AR, TB, SBor, AB, VFAM.

## Data and materials availability

All data and code used in the analysis are available on GitLab (https://git.scicore.unibas.ch/TBRU/mtuberculosis_epistasis) and the initial proteome identification and quantification file is available from Zenodo (https://doi.org/10.5281/zenodo.16744470).

## Supplementary Figures and Tables

**Fig. S1.**
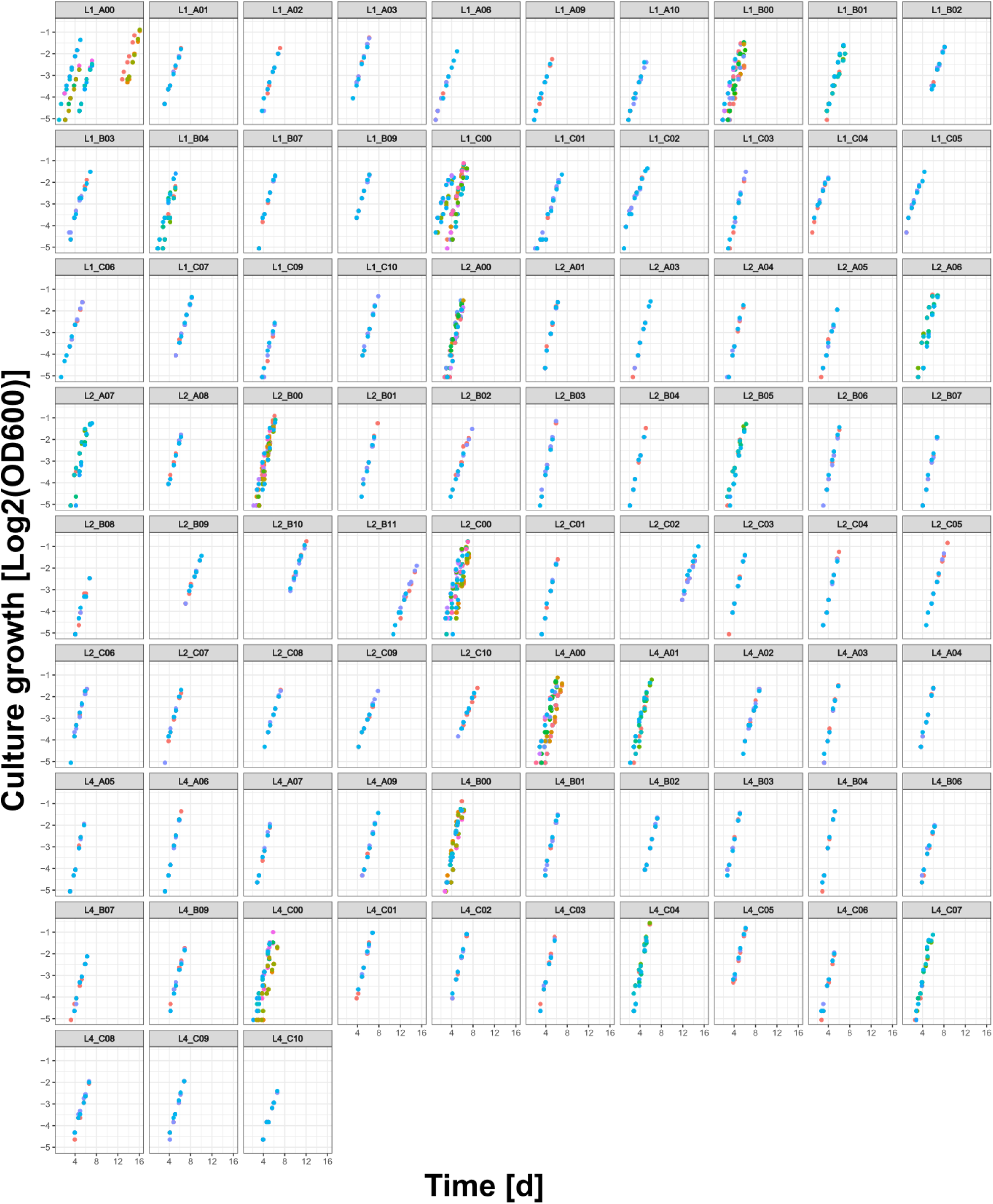
Per-strain single-replicate raw growth data after filtering for exponential phase (definition see methods). Each color is a different replicate. Strains with more than three replicates were measured in multiple experiments.

**Fig. S2.**
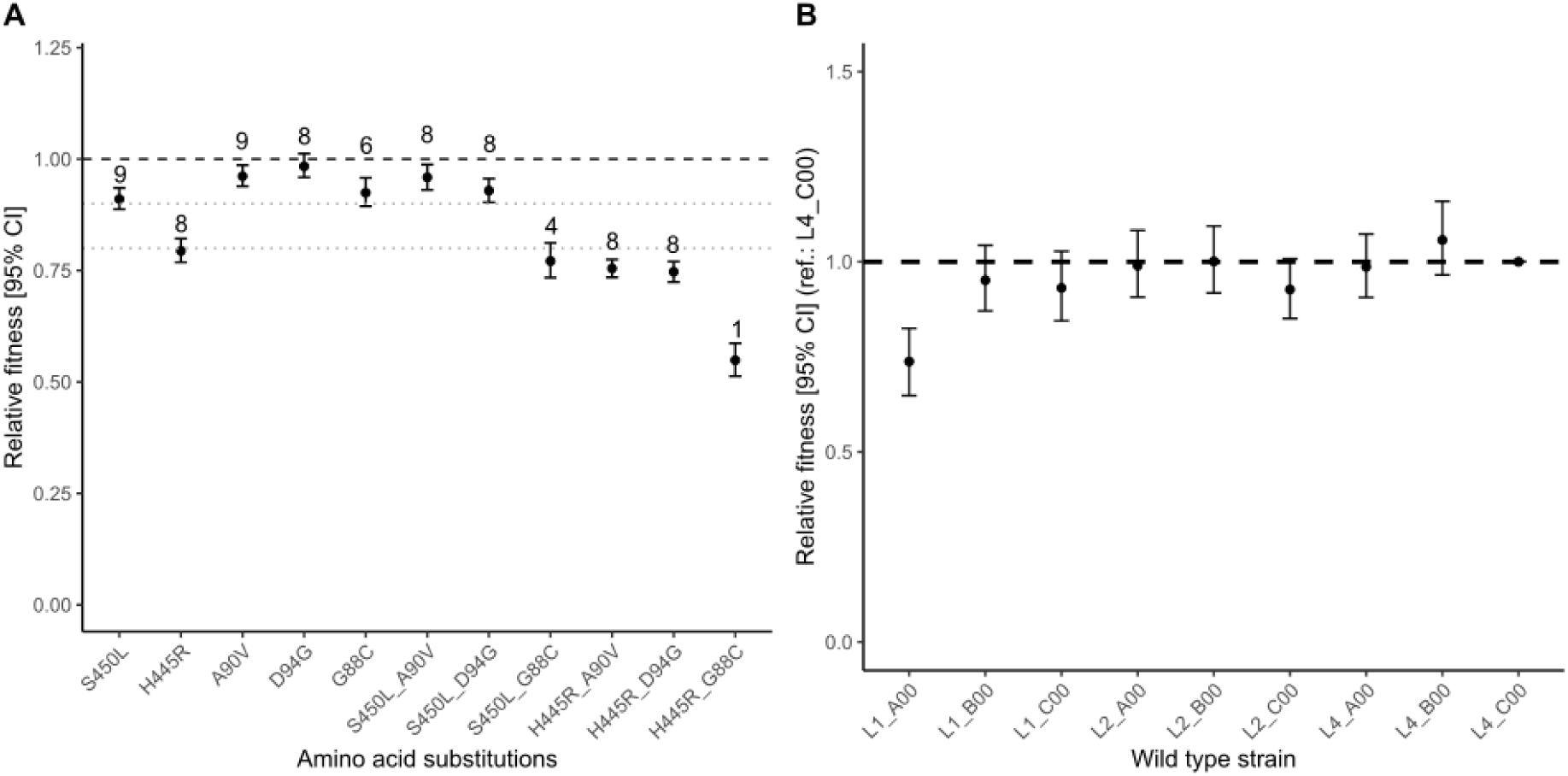
Relative fitness of drug resistance mutations and wild type strains. **A)** Relative fitness of mutants sharing specific RpoB and/or GyrA drug resistance mutations, across strain- backgrounds. Black dashed line = relative fitness = 1, gray dotted lines at relative fitness 0.9 and 0.8. Numbers of strains per mutation(-combination) indicated above error bars. **B)** Relative fitness of wild type strains compared to wild type strain L4_C00.

**Fig. S3.**
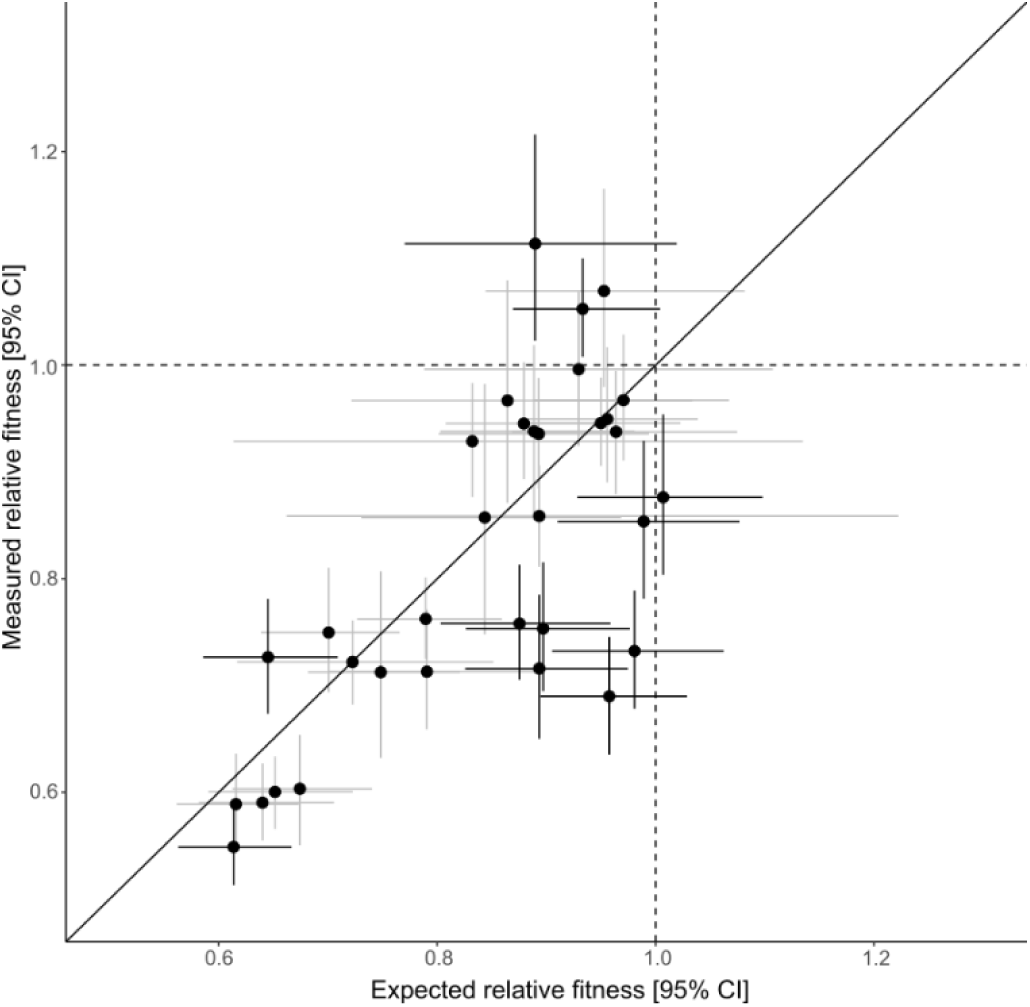
Evidence for epistasis between RpoB and GyrA drug resistance mutations. The measured vs. the expected (based on the relative fitness of the single-mutants assuming a multiplicative model) relative fitness of the double-mutants. 95% CIs with evidence of epistasis (n=11) are shown in black and with no evidence of epistasis (n=22) in gray.

**Fig. S4.**
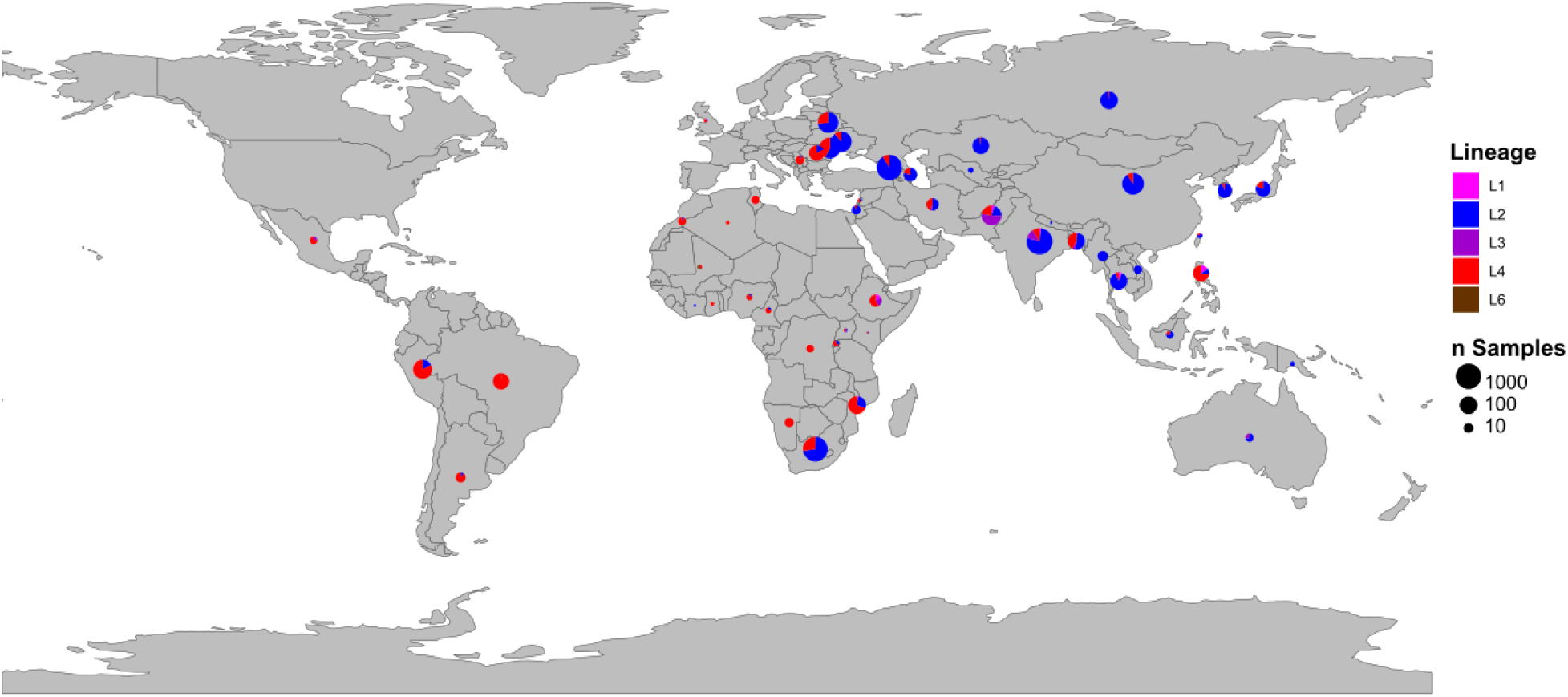
Global- and lineage-distribution of pre-XDR+ genomes. Of the 8,067 pre-XDR and XDR genomes, we could identify the origin of 7,517 samples, mapped here geographically.

**Fig. S5.**
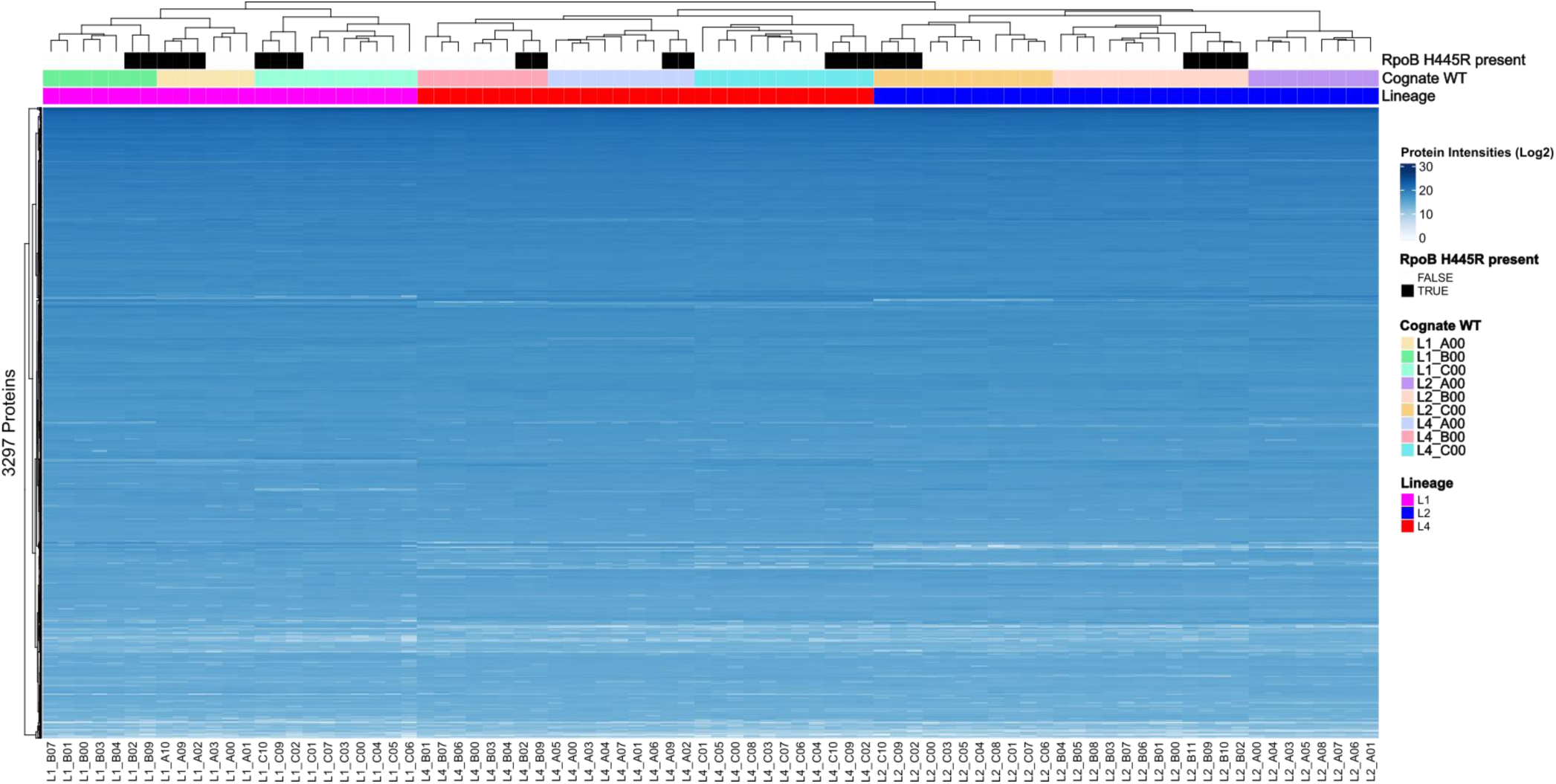
Heatmap of all proteome samples. Clustering occurred based on the log2-transformed protein intensities of 3,297 proteins. Black dashed lines = lineage-level separation, colored dashed lines = strain-level separation, red branches and labels = single- and double mutants with RpoB H445R

**Table S1.**
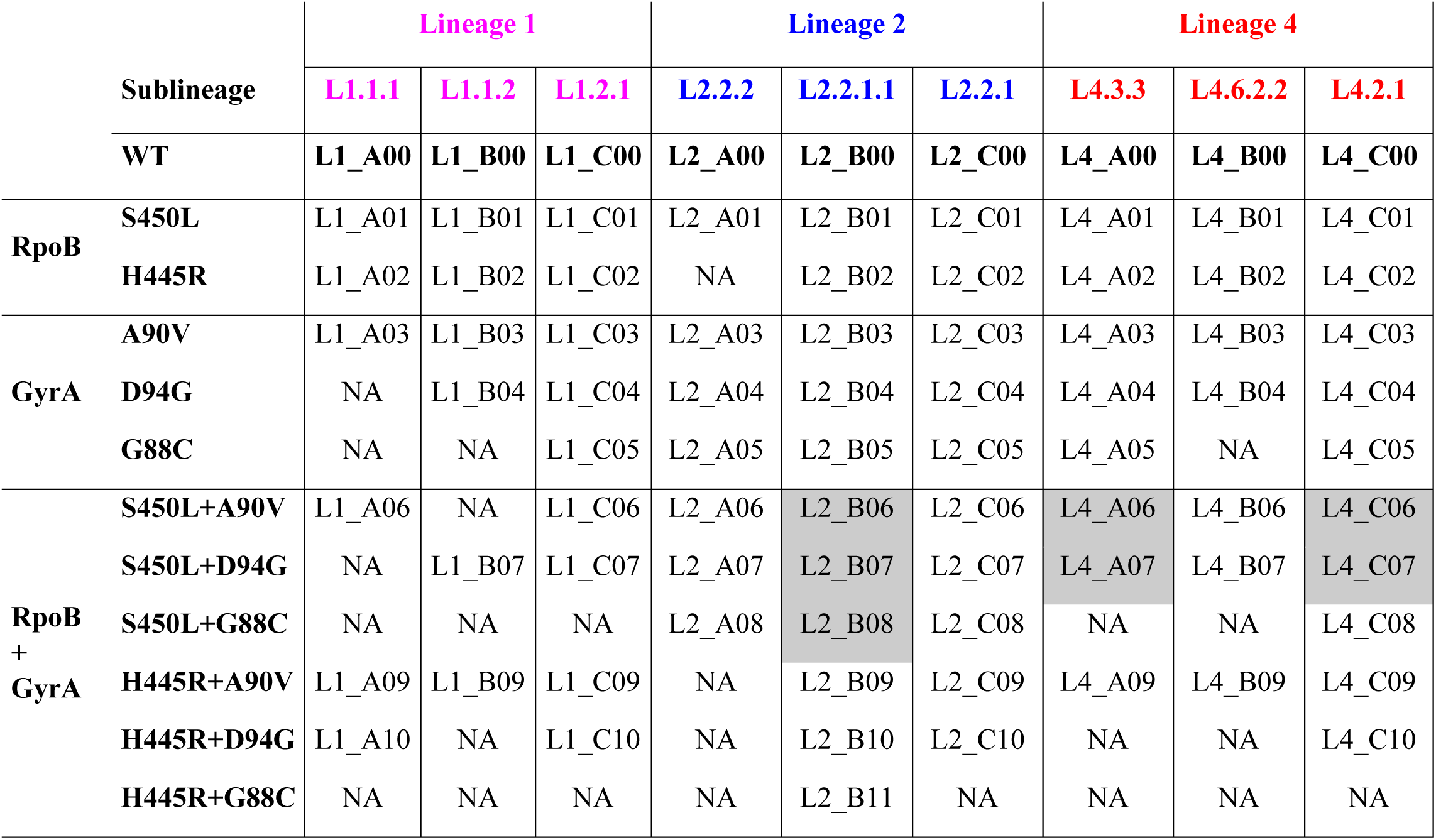
Wild type strains and their derived RpoB- and GyrA single- and double-mutants. NA = mutants we did not manage to obtain; gray boxes = strains with (potential) compensatory mutations.

**Table S2.** Nonsynonymous fixed off-target mutations.

| gene | mutation | strains | n | variant_category |
| --- | --- | --- | --- | --- |
| rpoA_Rv3457c | Thr181Ala | L2_B08; L2_B07; L2_B06 | 3 | Potential_compensatory |
| tgs3_Rv3234c | Glu16Lys | L1_B01; L1_B07 | 2 | NA |
| moaA1_Rv3109 | Ile136Val | L4_B02; L4_B09 | 2 | NA |
| ppk1_Rv2984 | Arg304Gln | L1_B02; L1_B09 | 2 | NA |
| rpoC_Rv0668 | Pro434Leu | L4_C07; L4_C06 | 2 | Compensatory |
| fadD28_Rv2941 | Arg288Pro | L4_C03 | 1 | NA |
| Rv3273_Rv3273 | Val707Leu | L4_C05 | 1 | NA |
| guaB3_Rv3410c | Gly273Arg | L4_C08 | 1 | NA |
| mbtE_Rv2380c | Ala422Thr | L2_A04 | 1 | NA |
| rpsK_Rv3459c | Lys83Thr | L1_B04 | 1 | NA |
| Rv3719_Rv3719 | Ile266Val | L4_C06 | 1 | NA |
| ppk1_Rv2984 | Arg316Gln | L1_A09 | 1 | NA |
| hrcA_Rv2374c | Val83Phe | L1_A10 | 1 | NA |
| rpoA_Rv3457c | Gly31Cys | L4_A06 | 1 | Potential_compensatory |
| wbbL1_Rv3265c | Pro26Gln | L2_C10 | 1 | NA |
| rpoA_Rv3457c | Val183Gly | L4_A07 | 1 | Compensatory |

**Table S3.**
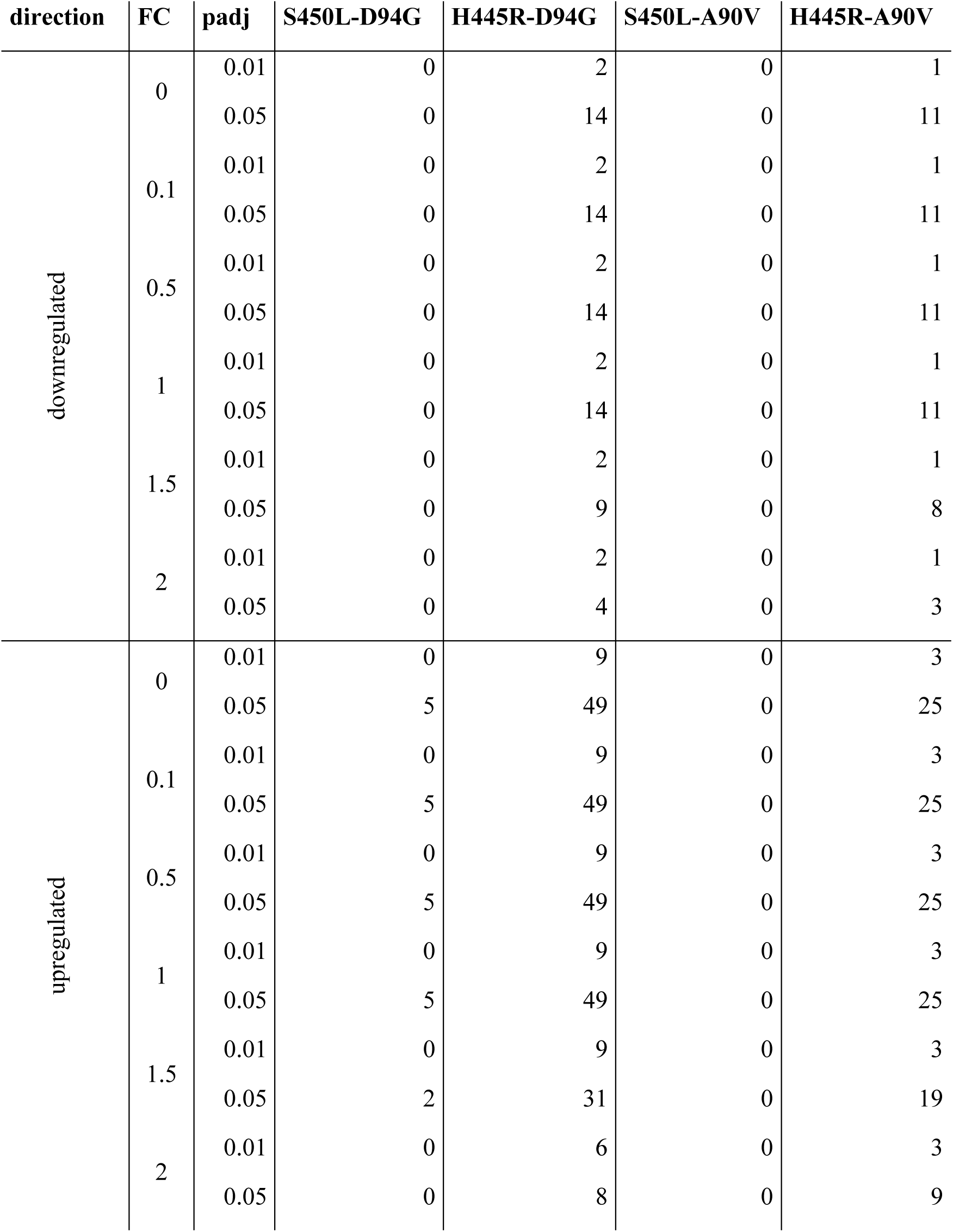
Sensitivity analysis with different thresholds for fold-change (FC) and adjusted p-value (padj). Only mutants of wild type strains, which have both, a S450L-D94G (n=4) and a H445R-D94G or a S450L-A90V and a H445R-A90V (n=6), were included in the analysis.

**Table S4.** Significant (p.adjust <0.01) results of enrichment analysis of significantly differentially abundant proteins in double mutants carrying RpoB S450L and H445R. Only mutants of wild type strains, which have both, a S450L-D94G and a H445R-D94G (n=4) or a S450L- A90V and a H445R-A90V (n=6), were included in the analysis. GR=GeneRatio, BR=BackgroundRatio, RF=RichFactor, FE=FoldEnrichment, n=number of significantly differentially abundant proteins identified in gene set, direction=direction of enrichment.

| ID | GR | BR | RF | FE | zScore | pvalue | p.adjust | qvalue | n | strain | RpoB | direction |
| --- | --- | --- | --- | --- | --- | --- | --- | --- | --- | --- | --- | --- |
| Aminoacyl-tRNA biosynthesis | 3/21 | 24/3973 | 0.12 | 23.65 | 8.11 | <0.001 | <0.01 | <0.01 | 3 | L1_C09 | H445R | down |
| Lipid metabolism | 14/69 | 270/3973 | 0.05 | 2.99 | 4.49 | <0.001 | <0.01 | <0.01 | 14 | L2_C09 | H445R | down |
| Mycolic acid biosynthesis | 6/69 | 36/3973 | 0.17 | 9.6 | 6.89 | <0.001 | <0.01 | <0.01 | 6 | L2_C09 | H445R | down |
| Lipid metabolism | 27/133 | 270/3973 | 0.1 | 2.99 | 6.29 | <0.001 | <0.001 | <0.001 | 27 | L2_C10 | H445R | down |
| Mycolic acid biosynthesis | 11/133 | 36/3973 | 0.31 | 9.13 | 9.12 | <0.001 | <0.001 | <0.001 | 11 | L2_C10 | H445R | down |
| Oxidative phosphorylation | 8/133 | 49/3973 | 0.16 | 4.88 | 5.08 | <0.001 | <0.01 | <0.01 | 8 | L2_C10 | H445R | down |
| 2-Oxocarboxylic acid metabolism | 4/32 | 35/3973 | 0.11 | 14.19 | 7.06 | <0.001 | <0.01 | <0.01 | 4 | L1_C10 | H445R | up |
| Pantothenate and CoA biosynthesis | 3/32 | 21/3973 | 0.14 | 17.74 | 6.93 | <0.001 | <0.01 | <0.01 | 3 | L1_C10 | H445R | up |
| Valine, leucine and isoleucine biosynthesis | 4/32 | 13/3973 | 0.31 | 38.2 | 12.11 | <0.001 | <0.001 | <0.001 | 4 | L1_C10 | H445R | up |
| Sulfur metabolism | 5/82 | 21/3973 | 0.24 | 11.54 | 7.03 | <0.001 | <0.01 | <0.01 | 5 | L4_A09 | H445R | up |
| Sulfur metabolism | 7/191 | 21/3973 | 0.33 | 6.93 | 6.13 | <0.001 | <0.01 | <0.01 | 7 | L4_C09 | H445R | up |
| Sulfur metabolism | 5/79 | 21/3973 | 0.24 | 11.97 | 7.18 | <0.001 | <0.01 | <0.01 | 5 | L4_C10 | H445R | up |
| PE/PPE | 3/3 | 165/3973 | 0.02 | 24.08 | 8.32 | <0.001 | <0.001 | <0.001 | 3 | L2_B06 | S450L | down |
| Ribosome | 6/62 | 56/3973 | 0.11 | 6.87 | 5.57 | <0.001 | <0.01 | <0.01 | 6 | L2_C06 | S450L | down |
| Information pathways | 21/120 | 241/3973 | 0.09 | 2.88 | 5.33 | <0.001 | <0.001 | <0.001 | 21 | L2_C07 | S450L | down |
| Ribosome | 10/120 | 56/3973 | 0.18 | 5.91 | 6.53 | <0.001 | <0.001 | <0.001 | 10 | L2_C07 | S450L | down |
| IdeR | 12/106 | 44/3973 | 0.27 | 10.22 | 10.18 | <0.001 | <0.001 | <0.001 | 12 | L4_B06 | S450L | down |
| Information pathways | 17/106 | 241/3973 | 0.07 | 2.64 | 4.36 | <0.001 | <0.01 | <0.01 | 17 | L4_B06 | S450L | down |
| Ribosome | 9/106 | 56/3973 | 0.16 | 6.02 | 6.27 | <0.001 | <0.001 | <0.001 | 9 | L4_B06 | S450L | down |

## References

1. H. J. Cordell, Epistasis: what it means, what it doesn’t mean, and statistical methods to detect it in humans. Hum Mol Genet 11, 2463–2468 (2002).

2. S. Trindade, A. Sousa, K. B. Xavier, F. Dionisio, M. G. Ferreira, I. Gordo, Positive epistasis drives the acquisition of multidrug resistance. PLoS Genet 5, e1000578 (2009).

3. S. Borrell, Y. Teo, F. Giardina, E. M. Streicher, M. Klopper, J. Feldmann, B. Muller, T. C. Victor, S. Gagneux, Epistasis between antibiotic resistance mutations drives the evolution of extensively drug-resistant tuberculosis. Evol Med Public Health 2013, 65–74 (2013).

4. H. Ward, G. G. Perron, R. C. Maclean, The cost of multiple drug resistance in Pseudomonas aeruginosa. J Evol Biol 22, 997–1003 (2009).

5. A. R. Hall, R. C. MacLean, Epistasis buffers the fitness effects of rifampicin-resistance mutations in Pseudomonas aeruginosa. Evolution 65, 2370–2379 (2011).

6. Geneva: World Health Organization, Global tuberculosis report 2024 (Geneva: World Health Organization, 2024).

7. Geneva: World Health Organization, WHO consolidated guidelines on tuberculosis. Module 4: treatment and care (Geneva: World Health Organization, 2025)

8. O. S. Pedersen, F. B. Holmgaard, M. K. D. Mikkelsen, C. Lange, G. Sotgiu, T. Lillebaek, A. B. Andersen, C. M. Wejse, V. N. Dahl, Global treatment outcomes of extensively drug-resistant tuberculosis in adults: A systematic review and meta-analysis. J Infect 87, 177–189 (2023).

9. C. Stritt, S. Gagneux, How do monomorphic bacteria evolve? The Mycobacterium tuberculosis complex and the awkward population genetics of extreme clonality. Peer Community Journal 3, (2023).

10. V. Eldholm, G. Norheim, B. von der Lippe, W. Kinander, U. R. Dahle, D. A. Caugant, T. Mannsaker, A. T. Mengshoel, A. M. Dyrhol-Riise, F. Balloux, Evolution of extensively drug-resistant Mycobacterium tuberculosis from a susceptible ancestor in a single patient. Genome Biol 15, 490 (2014).

11. I. Comas, S. Borrell, A. Roetzer, G. Rose, B. Malla, M. Kato-Maeda, J. Galagan, S. Niemann, S. Gagneux, Whole-genome sequencing of rifampicin-resistant Mycobacterium tuberculosis strains identifies compensatory mutations in RNA polymerase genes. Nat Genet 44, 106–110 (2011).

12. C. Loiseau, E. M. Windels, S. M. Gygli, L. Jugheli, N. Maghradze, D. Brites, A. Ross, G. Goig, M. Reinhard, S. Borrell, A. Trauner, A. Dotsch, R. Aspindzelashvili, R. Denes, K. Reither, C. Beisel, N. Tukvadze, Z. Avaliani, T. Stadler, S. Gagneux, The relative transmission fitness of multidrug-resistant Mycobacterium tuberculosis in a drug resistance hotspot. Nat Commun 14, 1988 (2023).

13. M. Merker, M. Barbier, H. Cox, J. P. Rasigade, S. Feuerriegel, T. A. Kohl, R. Diel, S. Borrell, S. Gagneux, V. Nikolayevskyy, S. Andres, U. Nubel, P. Supply, T. Wirth, S. Niemann, Compensatory evolution drives multidrug-resistant tuberculosis in Central Asia. Elife 7, (2018).

14. M. Merker, J. P. Rasigade, M. Barbier, H. Cox, S. Feuerriegel, T. A. Kohl, E. Shitikov, K. Klaos, C. Gaudin, R. Antoine, R. Diel, S. Borrell, S. Gagneux, V. Nikolayevskyy, S. Andres, V. Crudu, P. Supply, S. Niemann, T. Wirth, Transcontinental spread and evolution of Mycobacterium tuberculosis W148 European/Russian clade toward extensively drug resistant tuberculosis. Nat Commun 13, 5105 (2022).

15. G. A. Goig, F. Menardo, Z. Salaam-Dreyer, A. Dippenaar, E. M. Streicher, J. Daniels, A. Reuter, S. Borrell, M. Reinhard, A. Doetsch, C. Beisel, R. M. Warren, H. Cox, S. Gagneux, Effect of compensatory evolution in the emergence and transmission of rifampicin-resistant Mycobacterium tuberculosis in Cape Town, South Africa: a genomic epidemiology study. Lancet Microbe 4, e506–e515 (2023).

16. S. Borrell, A. Trauner, D. Brites, L. Rigouts, C. Loiseau, M. Coscolla, S. Niemann, B. De Jong, D. Yeboah-Manu, M. Kato-Maeda, J. Feldmann, M. Reinhard, C. Beisel, S. Gagneux, Reference set of Mycobacterium tuberculosis clinical strains: A tool for research and product development. PLoS One 14, e0214088 (2019).

17. D. A. Mitchison, J. B. Selkon, J. Lloyd, Virulence in the Guinea-Pig, Susceptibility to Hydrogen Peroxide, and Catalase Activity of Isoniazid-Sensitive Tubercle Bacilli from South Indian and British Patients. J Pathol Bacteriol 86, 377–386 (1963).

18. N. Krishnan, W. Malaga, P. Constant, M. Caws, T. H. Tran, J. Salmons, T. N. Nguyen, D. B. Nguyen, M. Daffe, D. B. Young, B. D. Robertson, C. Guilhot, G. E. Thwaites, Mycobacterium tuberculosis lineage influences innate immune response and virulence and is associated with distinct cell envelope lipid profiles. PLoS One 6, e23870 (2011).

19. M. Zwyer, L. K. Rutaihwa, E. Windels, J. Hella, F. Menardo, M. Sasamalo, G. Sommer, L. Schmulling, S. Borrell, M. Reinhard, A. Dotsch, H. Hiza, C. Stritt, G. Sikalengo, L. Fenner, B. C. De Jong, M. Kato-Maeda, L. Jugheli, J. D. Ernst, S. Niemann, L. Jeljeli, M. Ballif, M. Egger, N. Rakotosamimanana, D. Yeboah-Manu, P. Asare, B. Malla, H. Y. Dou, N. Zetola, R. J. Wilkinson, H. Cox, E. J. Carter, J. Gnokoro, M. Yotebieng, E. Gotuzzo, A. Abimiku, A. Avihingsanon, Z. M. Xu, J. Fellay, D. Portevin, K. Reither, T. Stadler, S. Gagneux, D. Brites, Back-to-Africa introductions of Mycobacterium tuberculosis as the main cause of tuberculosis in Dar es Salaam, Tanzania. PLoS Pathog 19, e1010893 (2023).

20. K. E. Holt, P. McAdam, P. V. K. Thai, N. T. T. Thuong, D. T. M. Ha, N. N. Lan, N. H. Lan, N. T. Q. Nhu, H. T. Hai, V. T. N. Ha, G. Thwaites, D. J. Edwards, A. P. Nath, K. Pham, D. B. Ascher, J. Farrar, C. C. Khor, Y. Y. Teo, M. Inouye, M. Caws, S. J. Dunstan, Frequent transmission of the Mycobacterium tuberculosis Beijing lineage and positive selection for the EsxW Beijing variant in Vietnam. Nat Genet 50, 849–856 (2018).

21. T. M. Walker, M. Choisy, M. Dedicoat, P. G. Drennan, D. Wyllie, F. Yang-Turner, D. W. Crook, E. R. Robinson, A. S. Walker, E. G. Smith, T. E. A. Peto, Mycobacterium tuberculosis transmission in Birmingham, UK, 2009-19: An observational study. Lancet Reg Health Eur 17, 100361 (2022).

22. L. Freschi, R. Vargas, Jr., A. Husain, S. M. M. Kamal, A. Skrahina, S. Tahseen, N. Ismail, A. Barbova, S. Niemann, D. M. Cirillo, A. S. Dean, M. Zignol, M. R. Farhat, Population structure, biogeography and transmissibility of Mycobacterium tuberculosis. Nat Commun 12, 6099 (2021).

23. K. A. Cohen, T. Abeel, A. Manson McGuire, C. A. Desjardins, V. Munsamy, T. P. Shea, B. J. Walker, N. Bantubani, D. V. Almeida, L. Alvarado, S. B. Chapman, N. R. Mvelase, E. Y. Duffy, M. G. Fitzgerald, P. Govender, S. Gujja, S. Hamilton, C. Howarth, J. D. Larimer, K. Maharaj, M. D. Pearson, M. E. Priest, Q. Zeng, N. Padayatchi, J. Grosset, S. K. Young, J. Wortman, K. P. Mlisana, M. R. O’Donnell, B. W. Birren, W. R. Bishai, A. S. Pym, A. M. Earl, Evolution of Extensively Drug-Resistant Tuberculosis over Four Decades: Whole Genome Sequencing and Dating Analysis of Mycobacterium tuberculosis Isolates from KwaZulu-Natal. PLoS Med 12, e1001880 (2015).

24. M. Merker, C. Blin, S. Mona, N. Duforet-Frebourg, S. Lecher, E. Willery, M. G. Blum, S. Rusch-Gerdes, I. Mokrousov, E. Aleksic, C. Allix-Beguec, A. Antierens, E. Augustynowicz-Kopec, M. Ballif, F. Barletta, H. P. Beck, C. E. Barry, 3rd, M. Bonnet, E. Borroni, I. Campos-Herrero, D. Cirillo, H. Cox, S. Crowe, V. Crudu, R. Diel, F. Drobniewski, M. Fauville-Dufaux, S. Gagneux, S. Ghebremichael, M. Hanekom, S. Hoffner, W. W. Jiao, S. Kalon, T. A. Kohl, I. Kontsevaya, T. Lillebaek, S. Maeda, V. Nikolayevskyy, M. Rasmussen, N. Rastogi, S. Samper, E. Sanchez-Padilla, B. Savic, I. C. Shamputa, A. Shen, L. H. Sng, P. Stakenas, K. Toit, F. Varaine, D. Vukovic, C. Wahl, R. Warren, P. Supply, S. Niemann, T. Wirth, Evolutionary history and global spread of the Mycobacterium tuberculosis Beijing lineage. Nat Genet 47, 242–249 (2015).

25. S. Borrell, S. Gagneux, Infectiousness, reproductive fitness and evolution of drug-resistant Mycobacterium tuberculosis. Int J Tuberc Lung Dis 13, 1456–1466 (2009).

26. Geneva: World Health Organization, Catalogue of mutations in Mycobacterium tuberculosis complex and their association with drug resistance (Geneva: World Health Organization, 2023)

27. R. A. D. Castro, A. Ross, L. Kamwela, M. Reinhard, C. Loiseau, J. Feldmann, S. Borrell, A. Trauner, S. Gagneux, The Genetic Background Modulates the Evolution of Fluoroquinolone-Resistance in Mycobacterium tuberculosis. Mol Biol Evol 37, 195–207 (2020).

28. S. Gagneux, C. D. Long, P. M. Small, T. Van, G. K. Schoolnik, B. J. Bohannan, The competitive cost of antibiotic resistance in Mycobacterium tuberculosis. Science 312, 1944–1946 (2006).

29. D. M. Weinreich, Watson, Richard A. and Chao, Lin, Perspective: Sign Epistasis and Genetic Constraint on Evolutionary Trajectories Evolution 59, 1165–1174 (2005).

30. K. E. Wiens, L. P. Woyczynski, J. R. Ledesma, J. M. Ross, R. Zenteno-Cuevas, A. Goodridge, I. Ullah, B. Mathema, J. F. Djoba Siawaya, M. H. Biehl, S. E. Ray, N. V. Bhattacharjee, N. J. Henry, R. C. Reiner, Jr., H. H. Kyu, C. J. L. Murray, S. I. Hay, Global variation in bacterial strains that cause tuberculosis disease: a systematic review and meta-analysis. BMC Med 16, 196 (2018).

31. G. A. Goig, E. M. Windels, C. Loiseau, C. Stritt, L. Biru, S. Borrell, D. Brites, S. Gagneux, Ecology, global diversity and evolutionary mechanisms in the Mycobacterium tuberculosis complex. Nat Rev Microbiol, (2025).

32. M. Zignol, A. S. Dean, N. Alikhanova, S. Andres, A. M. Cabibbe, D. M. Cirillo, A. Dadu, A. Dreyer, M. Driesen, C. Gilpin, R. Hasan, Z. Hasan, S. Hoffner, A. Husain, A. Hussain, N. Ismail, M. Kamal, M. Mansjo, L. Mvusi, S. Niemann, S. V. Omar, E. Qadeer, L. Rigouts, S. Ruesch-Gerdes, M. Schito, M. Seyfaddinova, A. Skrahina, S. Tahseen, W. A. Wells, Y. D. Mukadi, M. Kimerling, K. Floyd, K. Weyer, M. C. Raviglione, Population-based resistance of Mycobacterium tuberculosis isolates to pyrazinamide and fluoroquinolones: results from a multicountry surveillance project. Lancet Infect Dis 16, 1185–1192 (2016).

33. P. Xu, X. Li, M. Zhao, X. Gui, K. DeRiemer, S. Gagneux, J. Mei, Q. Gao, Prevalence of fluoroquinolone resistance among tuberculosis patients in Shanghai, China. Antimicrob Agents Chemother 53, 3170–3172 (2009).

34. Y. Che, Q. Song, T. Yang, G. Ping, M. Yu, Fluoroquinolone resistance in multidrug-resistant Mycobacterium tuberculosis independent of fluoroquinolone use. Eur Respir J 50, (2017).

35. A. Trauner, A. Banaei-Esfahani, S. M. Gygli, P. Warmer, J. Feldmann, M. Zampieri, S. Borrell, B. C. Collins, C. Beisel, R. Aebersold, S. Gagneux, Expression Dysregulation as a Mediator of Fitness Costs in Antibiotic Resistance. Antimicrob Agents Chemother 65, e0050421 (2021).

36. I. K. Jordan, I. B. Rogozin, Y. I. Wolf, E. V. Koonin, Essential genes are more evolutionarily conserved than are nonessential genes in bacteria. Genome Res 12, 962–968 (2002).

37. M. A. DeJesus, E. R. Gerrick, W. Xu, S. W. Park, J. E. Long, C. C. Boutte, E. J. Rubin, D. Schnappinger, S. Ehrt, S. M. Fortune, C. M. Sassetti, T. R. Ioerger, Comprehensive Essentiality Analysis of the Mycobacterium tuberculosis Genome via Saturating Transposon Mutagenesis. mBio 8, (2017).

38. K. Okonechnikov, O. Golosova, M. Fursov, U. team, Unipro UGENE: a unified bioinformatics toolkit. Bioinformatics 28, 1166–1167 (2012).

39. J. T. Belisle, M. G. Sonnenberg, Isolation of genomic DNA from mycobacteria. Methods Mol Biol 101, 31–44 (1998).

40. Stan Development Team, Stan Reference Manual, (2024).

41. R Core Team. R: A language and environment for statistical computing, (2024).

42. Stan Development Team, RStan: the R interface to Stan, (2024).

43. R. C. Lewontin, K.-i. Kojima, The Evolutionary Dynamics of Complex Polymorphisms. Evolution 14, (1960).

44. P. Massicotte, A. South. (2025).

45. M. Choi, C. Y. Chang, T. Clough, D. Broudy, T. Killeen, B. MacLean, O. Vitek, MSstats: an R package for statistical analysis of quantitative mass spectrometry-based proteomic experiments. Bioinformatics 30, 2524–2526 (2014).

46. J. P. Quast, D. Schuster, P. Picotti, protti: an R package for comprehensive data analysis of peptide- and protein-centric bottom-up proteomics data. Bioinform Adv 2, vbab041 (2022).

47. A. Banaei-Esfahani, A. Trauner, S. Borrell, S. M. Gygli, T. R. Rustad, J. Feldmann, L. C. Gillet, O. T. Schubert, D. R. Sherman, C. Beisel, S. Gagneux, R. Aebersold, B. C. Collins, Network analysis identifies regulators of lineage-specific phenotypes in Mycobacterium tuberculosis. (2020).

48. G. Yu, L. G. Wang, Y. Han, Q. Y. He, clusterProfiler: an R package for comparing biological themes among gene clusters. OMICS 16, 284–287 (2012).

49. S. Dray, A.-B. Dufour, Theade4Package: Implementing the Duality Diagram for Ecologists. Journal of Statistical Software 22, (2007).

